# Should regional species loss be faster, or slower, than local loss? It depends on density-dependent rate of death

**DOI:** 10.1101/2024.04.05.588218

**Authors:** Petr Keil, Adam T. Clark, Vojtěch Barták, François Leroy

**Affiliations:** Department of Spatial Sciences, Faculty of Environmental Sciences, Czech University of Life Sciences Prague, Kamýcká 129, 16500 Praha-Suchdol, Czech Republic; Institute of Biology, University of Graz, Holteigasse 6, 8010 Graz, Austria

**Keywords:** species richness, extirpation, sixth mass extinction, scaling, community dynamics, metacommunity, mortality

## Abstract

Assessment of the rate of species loss, which we also label extinction, is an urgent task. However, the rate depends on spatial grain (average area *A*) over which it is assessed—local species loss can be on average faster, or slower, than regional or global loss. Ecological mechanisms behind this discrepancy are unclear. We propose that the relationship between extinction rate and *A* is driven by two classical ecological phenomena: the Allee effect and the Janzen-Connell effect. Specifically, we hypothesize that (i) when per-individual probability of death (*P*_*death*_) decreases with population density *N* (as in Allee effects), per-species extinction rate (*Px*) should be high at regional grains, and low locally. (ii) In contrast, when *P*_*death*_ increases with *N* (as in Janzen-Connell effects), *Px* should be low regionally, but high locally. (iii) Total counts of extinct species (*Ex*) should follow a more complex relationship with *A*, as they also depend on drivers of the species-area relationship (SAR) prior to extinctions, such as intraspecific aggregation, species pools, and species-abundance distributions. We tested these hypotheses using simulation experiments, the first based on point patterns, the second on a system of generalized Lotka-Volterra equations. In both experiments, we used a single continuous parameter that moved between the Allee effect, no relationship between *P*_*death*_ and *N*, and the Janzen-Connell effect. We found support for our hypotheses, but only when regional species-abundance distributions were uneven enough to provide sufficiently rare or common species for Allee or Janzen-Connell to act on. In all, we have theoretically demonstrated a mechanism behind different rates of biodiversity change at different spatial grains which has been observed in empirical data.

## 2 Introduction

Excessive loss of biodiversity via species extinctions and extirpations is a serious threat to human wellbeing and ecosystem functioning (IPBES 2019). Assessments of how fast and where species disappear are thus necessary to identify causes of the loss, and for effective conservation decisions. The problem with such assessments is that both species diversity, and its loss, strongly depend on the area over which they are assessed.

When biodiversity is measured across multiple locations, e.g. in a grid on a map, the average area of a location is *spatial grain* (hereafter *A*, Table 1). Average species diversity can only increase or remain constant with increasing grain (Arrhenius 1921, Storch 2016). However, temporal change of diversity, and particularly extinction rates, can increase, decrease, or can have complex and non-linear relationships with grain (Keil et al. 2018, Chase et al. 2019). Specifically, the average number of species that have disappeared from typical local patches (fine grain) may be higher, or lower, than the number of species that have disappeared from a typical large region (coarse grain). This has practical consequences: First, reports of biodiversity loss from a single grain can mask more, or less, dramatic losses at other grains. Second, analyses of the loss conducted at different grains are not comparable.

**Table 1.**
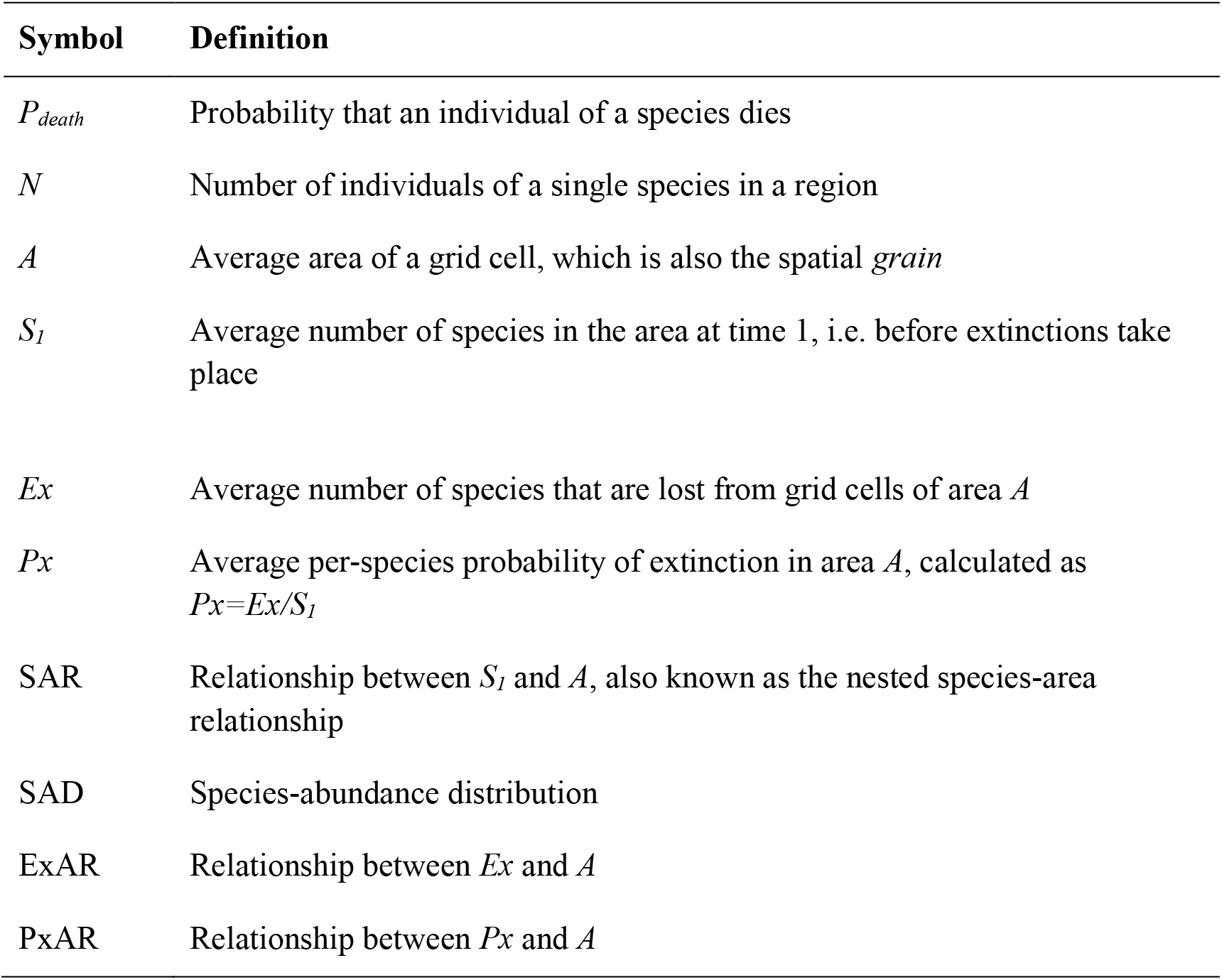
Key terms and notation used in this paper. When we mention averages, we mean that the value was averaged across all grid cells (or locations) at a given grain *A*.

Furthermore, the relationship between grain *A* and the rate of species loss depends on the metric of the loss. For example, the average number of species which went extinct (*Ex*, Table 1) can increase with increasing *A*, while the average per-species probability of extinction (*Px*, Table 1) can decrease with increasing *A*, with all of this happening in the same region and taxonomic group (Keil et al. 2018).

Mechanisms behind this strong and complex grain-dependency of diversity loss are poorly understood. Keil et al. (2018) showed that the grain dependence of extinction rates should be widely expected, but they provided only a simplistic cartoon of the mechanisms behind it (in their Fig. 2). Yan et al. (2022) further explored how the relationship between local and regional extinction rates depends on the spatial distribution of species, and they found that it is affected by spatial aggregation and mean range size. However, so far, we lack theory linking the shape of the extinction rate-area relationship with specific ecological processes on the individual level.

**Figure 1.**
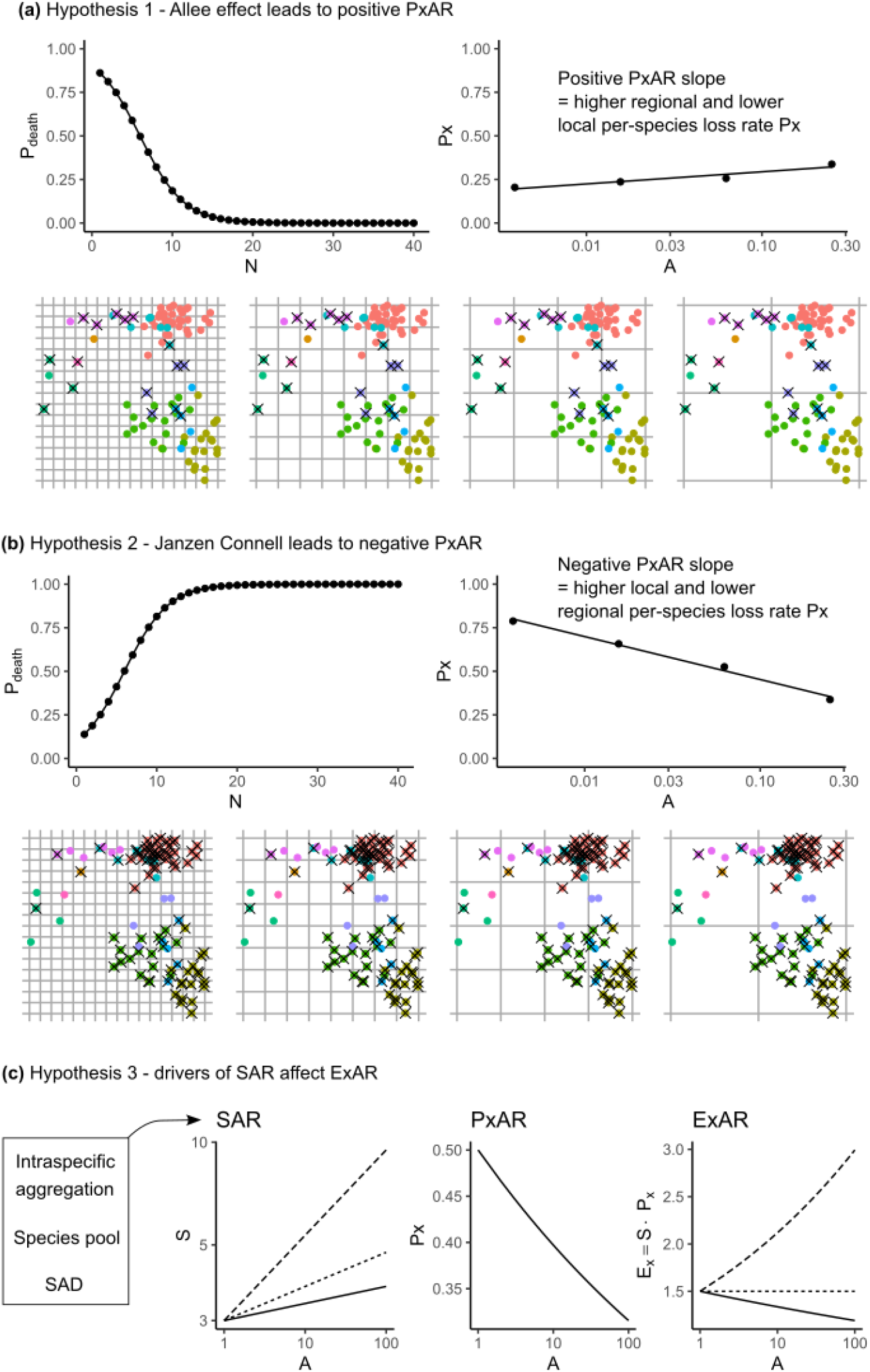
Three hypotheses of this study. Hypotheses 1 and 2 (a, b) predict different slopes of PxAR, which is the relationship between *per-species* probability of extinction (*Px*) and area (*A*). (a) Allee effect causes the *per-individual* probability of death (*P*_*death*_) to decrease with the total number of individuals in a region (*N*), leading to a positive slope of PxAR. (b) Janzen-Connell effect causes *P*_*death*_ to increase with *N*, leading to a negative slope of PxAR. Both hypotheses are illustrated on a point pattern in a square region, each individual is a dot, species are indicated by colors, and dead individuals are marked by a cross. The region is overlaid by four grids with increasing spatial grain *A*. (c) Hypothesis 3 states that the slope of ExAR, which is the relationship between number of extinct species (*Ex*) and *A*, is affected by drivers of the slope of species-area relationship (SAR) before extinctions take place. These drivers are intraspecific aggregation, species pool, and the shape of the regional species-abundance distribution (SAD).

**Figure 2.**
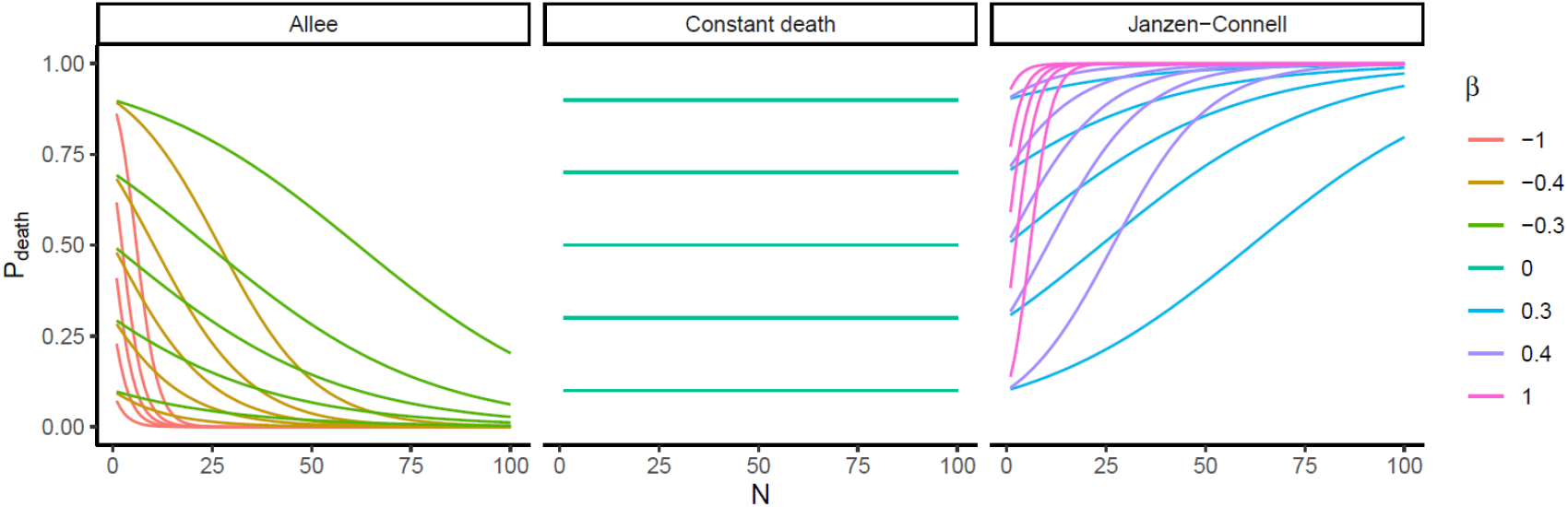
Illustration of how parameter *β* affects the shape of the Barták function (eq. 1). *β*<0 is the Allee effect (left panel), *β*= 0 is a density-independent *P*_*death*_ with constant values for all population sizes (middle panel), and *β*>0 is Janzen-Connell effect (right panel). The curves also vary in parameter *α*, which is the intercept of the Barták function.

In this paper we address this knowledge gap. We build on the idea in Fig. 2 of Keil et al. (2018) which suggests that the direction (positive, or negative) of the relationship between per-species extinction probability *Px* and grain *A* (we call this relationship *PxAR*; Table 1) somehow depends on species’ rarity. Here we propose (and test) that the key quantity is per-individual probability of death (*P*_*death*_), and particularly its relationship with population density of the species in the entire region. This relationship can either be positive or negative, which will then lead to negative or positive relationship between grain and extinction rate *Px* respectively. Here is why:

### Hypothesis 1

*If individuals of rare species are more likely to die than individuals of common species (as in Allee effect), there will be lower local and higher regional per-species probability of extinction Px (Fig. 1a)*. This hypothesis assumes a *negative* relationship between the number of individuals (*N*) in a region, and the per-individual probability of death (*P*_*death*_). As the number of individuals decreases, each of them is more likely to die (Fig. 1a). We consider this density-dependent death rate to arise from the Allee effect (Allee and Bowen 1932, Courchamp et al. 2009), which can occur, e.g., in organisms that rely on intraspecific facilitation (Berec et al. 2007), in organisms that alter their environment to suit them better, or in populations susceptible to inbreeding depression or demographic stochasticity (Lande et al. 2003). Consequently, species with fewer individuals are more likely to go extinct than species with more individuals. A loss of a rare small population reduces the total number of species in a region, but not so much the average number of species in a local patch. Thus, under the (hugely) simplifying assumption that the Allee effect applies to all species in a region, we expect local extinction rates to be lower than regional ones. In other words, the average *per-species* probability of extinction (*Px*, Table 1) should increase with grain *A* (Fig. 1a), i.e. PxAR should have a positive slope.

### Hypothesis 2

*If individuals of common species are more likely to die than individuals of rare species (as in Janzen-Connell effect), there will be higher local and lower regional per-species probability of extinction Px* (Fig. 1b). This hypothesis assumes a *positive* relationship between the number of individuals (*N*) in a region, and the per-individual probability of death (*P*_*death*_). The more individuals there are, the more likely each of them is to die. This is a simplified form of the Janzen-Connell effect (Connell 1970, Janzen 1970), which takes place when, e.g., large population densities lead to increases in the densities of pathogens or other natural enemies, or in intraspecific competition. Consequently, a decline in the population of common species (with high *N*) is unlikely to cause an extinction across the whole region (and once *N* declines to below a certain level, the effect no longer applies), but it should cause more frequent extinctions of the species from local patches, affecting local extinction rates more than regional ones. Thus, assuming the Janzen-Connell effect applies equally to all species in the community, the average per-species probability of extinction (*Px*) should decrease with area *A* (Fig. 1a), i.e. PxAR should have a negative slope.

### Hypothesis 3

*The relationship between the average counts of extinct species (Ex) and grain (we call this relationship ExAR; Table 1) should also be affected by Allee or Janzen-Connell effects. However, ExAR also depends on the initial number of species (S), and hence it should be sensitive to variables affecting the slope of the initial species-area relationship (SAR) in the region at time before extinctions take place, namely to species aggregation, the species pool, and the regional species-abundance distribution* (Fig. 1c). It is more difficult to predict how Allee and Janzen-Connell affect ExAR. This is because, since *E*_*x*_ = *S* ×*P*_*x*_ at a given grain, *Ex* is affected both by *Px* and by the number of species (*S*) present before the extinctions take place. Species richness *S* follows a nested SAR (Storch 2016), and thus ExAR= SAR×PxAR. Therefore, ExAR should be affected both by the Allee and Janzen-Connell effects (through PxAR, if hypotheses 1 and 2 hold), and by the slope of the *SAR*. The most important drivers of the SAR slope are *intra-specific spatial aggregation*, size of the *species pool*, and the shape of the regional *species-abundance distribution* (SAD) (Storch et al. 2008, McGlinn et al. 2019). We thus expect all these to affect the slope of the ExAR in addition to the Allee and Janzen-Connell effects. However, the interplay can be complex, as even a simple monotonic PxAR and SAR can lead to a plethora of functional forms of ExAR, including nonlinear or hump-shaped ExARs, as demonstrated by Keil et al. (2018).

## 3 Methods

We performed two simulation experiments (code and simulated data are available at https://github.com/petrkeil/extinction_scaling_simulations), and we tested all three hypotheses in each experiment. The first experiment is based on spatially explicit point patterns, it has two time steps (before and after the extinction), and hence it represents non-equilibrium community dynamics. The second experiment is based on a spatially implicit system of generalized Lotka-Volterra equations with many time steps and thus represents a community at equilibrium. As these two experiments are conceptually different, a result that emerges consistently in both can be seen as general, while differences among them can point to interesting ecological mechanisms.

### 3.1 Point pattern experiment

We first tested our hypotheses on a set of simulated point patterns; these have the advantage of being spatially explicit and allowed testing Hypothesis 3 under varying total numbers of species, individuals, shapes of the initial species-abundance distributions (SAD), and spatial aggregation of individuals, all of which affect the SAR (McGlinn et al. 2019). Spatial aggregation has also been shown to affect the spatial scaling of extinction rates (Yan et al. 2022). We did the simulations in R (R Development Core Team 2022) using a combination of the ‘mobsim’ package (May et al. 2018) and our custom functions (complete code is in the GitHub repository). Specific parameters of the simulations are in Table 2. Each simulation had 3 steps:

**Table 2.**
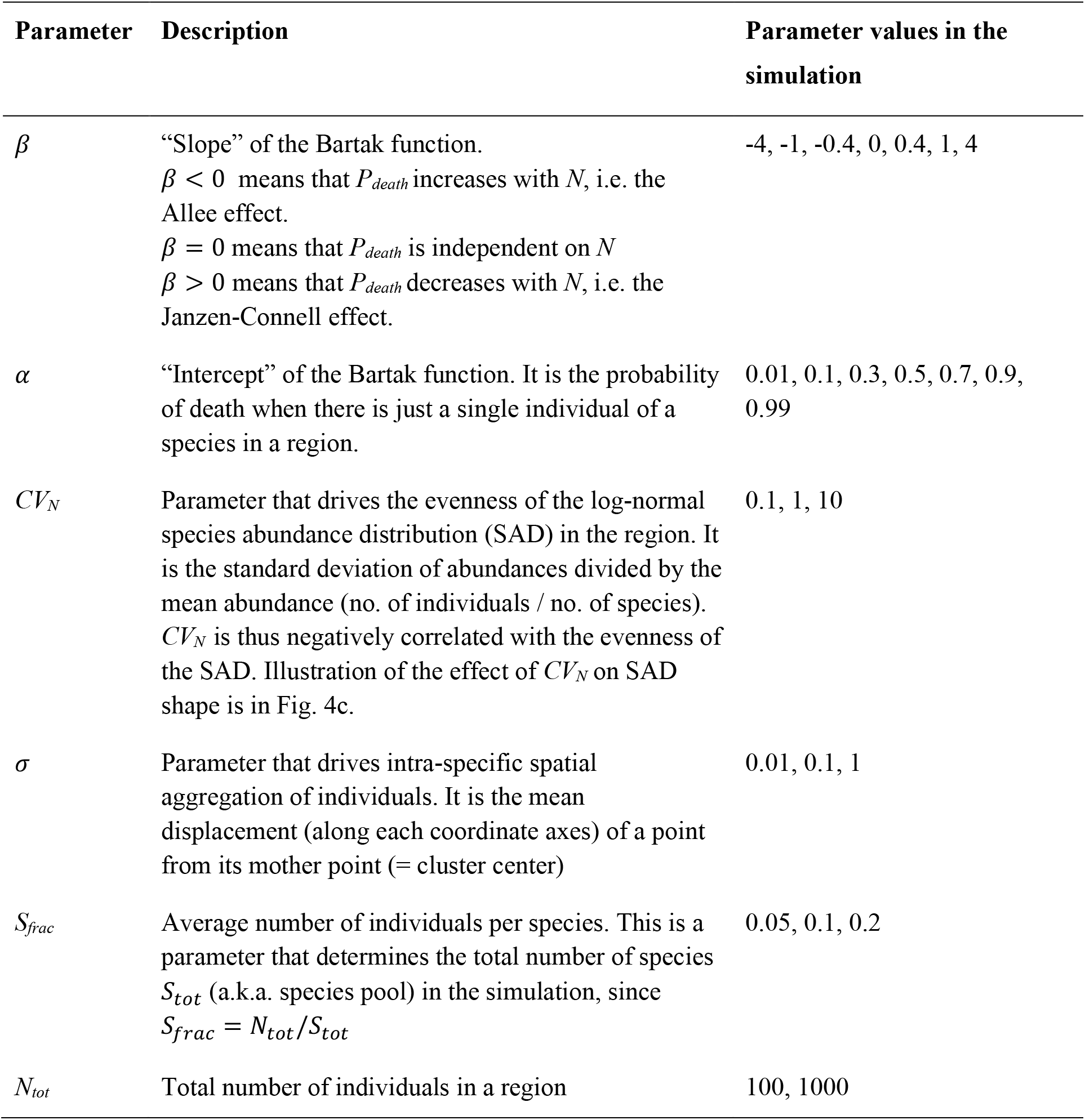
Parameters which we varied in the point pattern simulations.

#### 1. Species distributions in time 1

The simulations took place in a square region with an area of 1 imaginary unit (as shown in Fig. 1). We populated the region with a total number of *N*_*tot*_ points and *S*_*tot*_ species, using Thomas point process, each species with a single mother point and displacement from the point given by parameter *σ*. The number of individuals (*N*) of each species was given by a lognormal SAD whose evenness was set by a single parameter *CV*_*N*_ (explained in Table 2).

#### 2. Death of individuals

In the next step, we subjected every individual in the region to death with probability *P*_*death*_, which was made density-dependent according to a function:

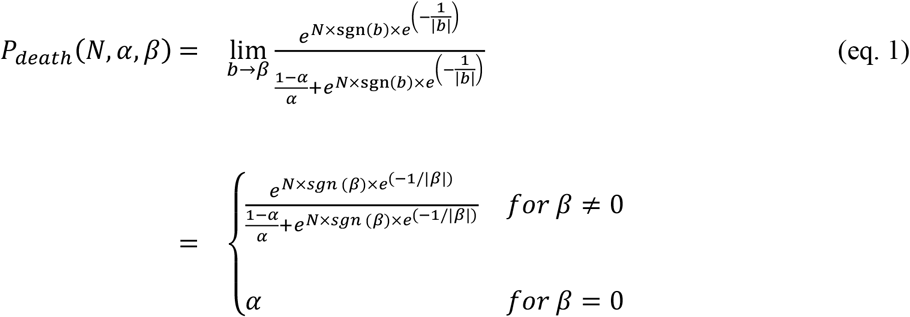

where *N* ∈{1,2, …}is the total number of individuals of a species in the region. We call eq. 1 the Barták function. Eq. 1 is a two-parameter function: (1) Parameter *α* ∈ (0, 1) is the “intercept”, i.e. the probability of an individual’s death when there is just a single individual in the region. (2) Parameter *β*∈ ℝ is the “slope” of the Barták function, where *β*<0 makes the function decreasing (Allee effect), *β*>0 makes it increasing (Janzen-Connell effect), and *β*= 0 means that there is no relationship between *N* and *P*_*death*_ (Fig. 2). Note that the Barták function has no biological nor mechanistical ground; it is a purely empirical function which enabled to continuously move between different magnitudes of the Allee and Janzen-Connell effects, using a single parameter *β*. In other words, this parameter *β*determines the strength (and sign) of the density dependence of the death rate. To execute the actual deaths of the individuals, we subjected each individual point to a Bernoulli trial with *P*_*death*_ as its *P* parameter.

#### 3. Calculation of species loss

In the final step, we overlaid the region with grids of increasing resolutions, where we divided the region into 2×2, 4×4, 8×8, and 16×16 grid cells. At each resolution we calculated *Ex* and *Px* (Table 1). We then fitted the PxAR as a linear regression of *Px* as a function of log(*A*), and ExAR as a Poisson generalized linear model (log link function) of *Ex* as a function of log(*A*).

We repeated the three steps above for 2,646 combinations of total number of individuals, size of species pool, shape of SAD, intra-specific spatial aggregation, and slope and intercept of the Barták function (Table 2). Each combination was repeated 10 times, giving us 26,460 simulation runs.

### 3.2 Lotka-Volterra experiment

To complement the point pattern simulations, we tested our hypotheses using a spatially implicit metacommunity simulation where abundances of species were traced in a set of *M* local communities, each following a disordered systems model (described below) comprising a series of generalized Lotka-Volterra equations (Barbier et al. 2018). Notably, the shape of the species-abundance distribution (SAD) was not specified à priori (as in experiment 1), but it emerged from the simulation. Thus, the magnitude of the density dependence of death rate (parameter *d*_*z*_, Table 3, similar to the *β* parameter for the point pattern simulations) affected the resulting SAD. We expected this to affect the resulting scaling: For example, if Allee effect drives *P*_*death*_, then rare species are constantly eliminated in each simulation step; over the long run, this should result in an even SADs with less rare species, and thus less extinctions, which could potentially “switch off” hypotheses 1 and 2.

**Table 3.**
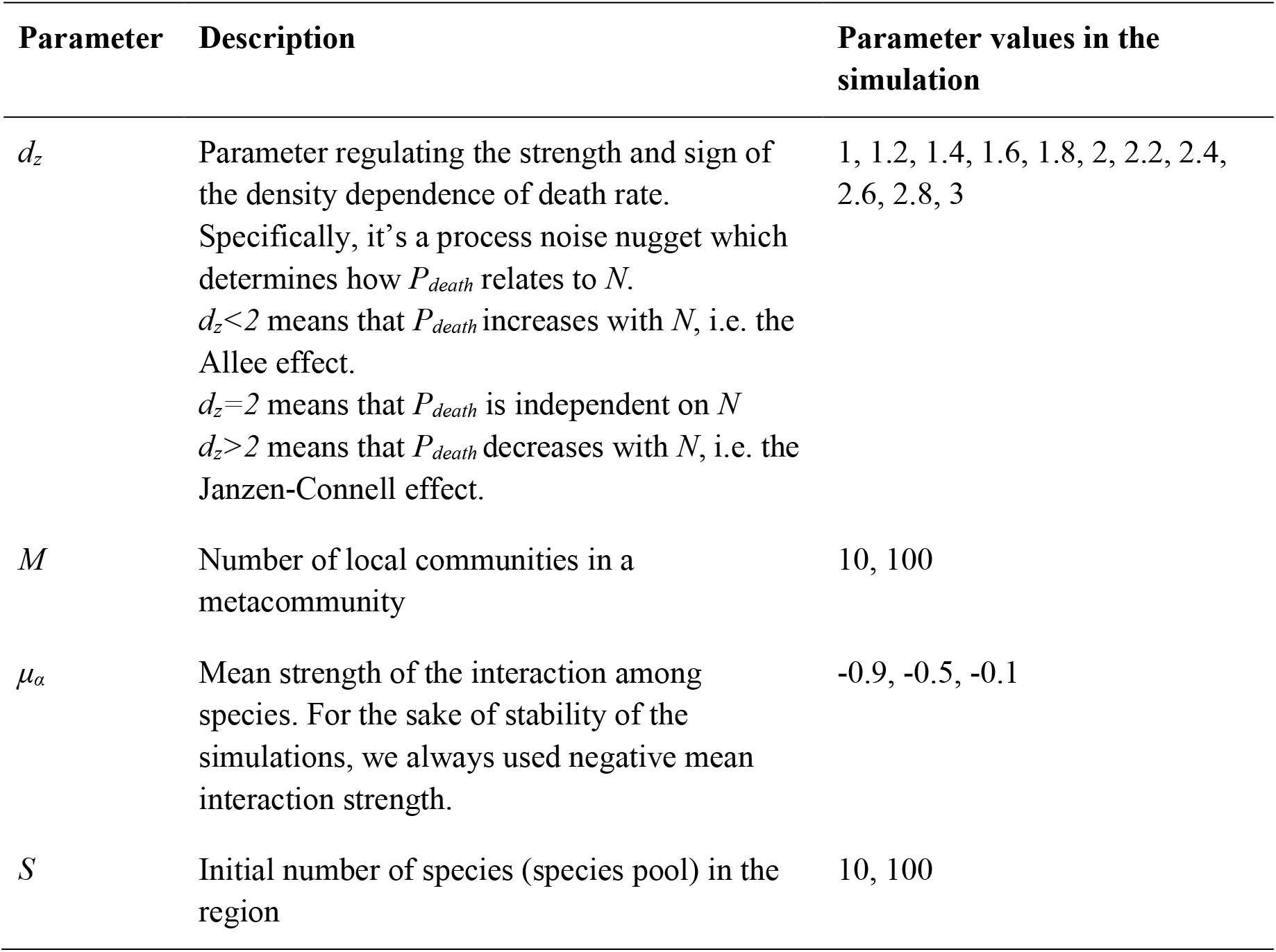
The key parameters which we varied in the Lotka-Volterra simulations. There are additional parameters which we kept constant, they are listed in the documented code’s repository at https://github.com/petrkeil/extinction_scaling_simulations.

The simulations proceeded as follows:

#### 1. Initial state

There are *M* patches, with a total of *S* species. Each species *i* has a carrying capacity (*K*_*i*_), per-capita growth rate (*r*_*i*_), and interspecific interaction strengths describing the per-capita effect of each species *j* on species *i* (*α*_*j,i*_). These are drawn from standard normal distributions, with means *μ*_*K*_, *μ*_*r*_, and *μ*_*α*_, and standard deviations *σ*_*K*_, *σ*_*r*_, and *σ*_*α*_, respectively (with minimum value 0 for *K*_*i*_ - note that *α*_*j,i*_ can be positive or negative). For simplicity, we assume that *α*_*j,i*_ and *α*_*i,j*_ are drawn independently.

#### 2. Dispersal

Species disperse across *M* patches, with dispersal and patch-level disturbance events modeled as a series of discrete stochastic perturbations to the underlying Lotka-Volterra models, with exponentially distributed waiting times between events. Each species in each patch disperses to new patches with rate *c*_*i*_ and initial abundance *n*_*0,i*_, drawn from normal distributions with mean *μ*_*c*_, and *μ*_*n0*_, standard deviations *σ*_*c*_ and *σ*_*n0*_, and minimum value 0. We set *n*_*0,i*_ of all species to the absolute value of the random draw.

#### 3. Extinction

Local extinction of a species within a local patch *m* (i.e. abundance of species *i, n*_*im*_(*t*) = 0) occurs either because of species interactions, or from impacts of disturbances. Disturbances impact all species and patches simultaneously, with the average waiting time between disturbances *d*_*w*_. Disturbance events are drawn from a normal distribution with mean *µd* = 0 and standard deviation 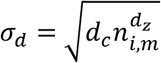, where *n*_*i,m*_ is the abundance of species *i* in patch *m*, and *d*_*c*_ and *d*_*z*_ are constants. Disturbances therefore impart stochastic structure to population dynamics that follow a Taylor Power Law (Taylor 1961). Thus, disturbance intensity grows as a nonlinear function of abundance, such that *d*_*z*_ < 2 implies that disturbances are more likely to drive rare species to local patch-level extinction, whereas *d*_*z*_ > 2 implies that common species are more likely to be driven to local extinction.

We tested all combinations of parameter values in Table 3, each repeated 10 times, giving us 1,320 simulations in total.

Each simulation was run for 20 steps, and we took a snapshot of species composition in all patches in step 10 and 20. A species was considered extinct in a patch (or in the whole metacommunity) if it was present in time 10 and absent in time 20. To calculate PxAR and ExAR, we calculated the average *Px*_*patch*_ and *Ex*_*patch*_ across all patches, and *Px*_*metacom*_ and *Ex*_*metacom*_ at the metacommunity scale. The slope of the PxAR was then (*Px*_*metacom*_ −*Px*_*patch*_)/*t* and the slope of ExAR was (*Ex*_*metacom*_ − E*x*_*patch*_)/*t*, where *t* = 10, which is the time between the two measurements.

#### Evaluating the simulations

For each simulation we noted the parameter values (Table 2, 3) as well as the slope of the PxAR and ExAR. We then plotted the slope of the PxAR and ExAR as a function of the parameter *β*(in point patterns simulations), or as a function of parameter *d*_*z*_ (in Lotka-Volterra simulations).

To compare the effects of different simulation parameters on the slope of PxAR and ExAR, we did a variable importance analysis using the Random Forest algorithm (Hastie et al. 2011) with the R package randomForest (Liaw and Wiener 2002) in which the slope of the PxAR or ExAR was the response variable, and parameters of the simulation (Table 2, 3) were the predictors. For the hyperparameters, we used the default settings of the randomForest R function.

We note that it makes little sense to evaluate importance of simulation parameters using P-values (White et al. 2014). We thus make our inference by directly comparing the frequency distributions of PxAR and ExAR slopes, or the variable importance from the random forest analyses.

## 4 Results

In both experiments, and among all other parameters considered, the parameters that controlled the magnitude of the Allee vs Janzen-Connell effects (*β* or *d*_*z*_) had the most important effect on slopes of both PxAR and ExAR (Fig. 3). In point pattern simulations, the single-individual *P*_death_ (i.e., the parameter *α*) was also relatively important. The slope of the ExAR was also relatively sensitive to the species aggregation (parameter *σ*) and the average species abundance (parameter *S*_fraq_).

**Figure 3.**
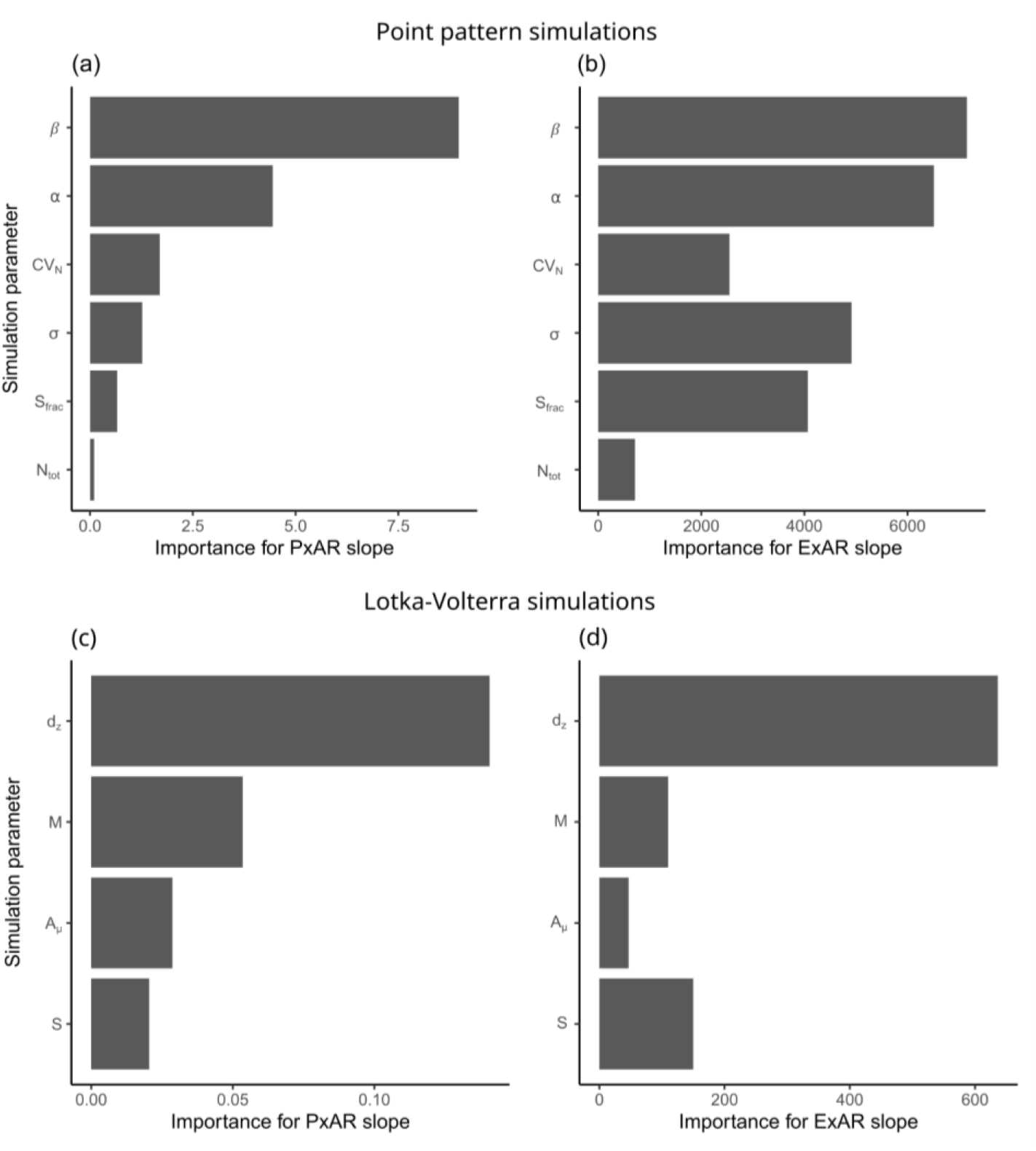
Importance of simulation parameters for determining the slope of PxAR and ExAR. For explanation of parameters see Tables 2 and 3. The importance is measured as the total decrease in node impurities from splitting on a given predictor variable, averaged over all trees in a random forest analysis.

### Hypothesis 1

We found that presence of the Allee effect in the point pattern simulations led to positive slopes of PxAR (Fig. 4a, red boxes), consistent with Hypothesis 1, but only when there were species with relatively low *N* in the regional SAD, i.e. when the *CV*_*N*_ was high and the SAD was uneven. For even SADs (*CV*_*N*_*=0*.*1*) the Allee effect produced a PxAR slope that was higher than a constant *P*_*death*_ scenario (Fig. 4a, green box), but often negative. In the Lotka-Volterra simulations, we didn’t find support for Hypothesis 1, although simulations with Allee effect were the only ones that produced the rare positive PxAR slopes (Fig. 5a).

**Figure 4.**
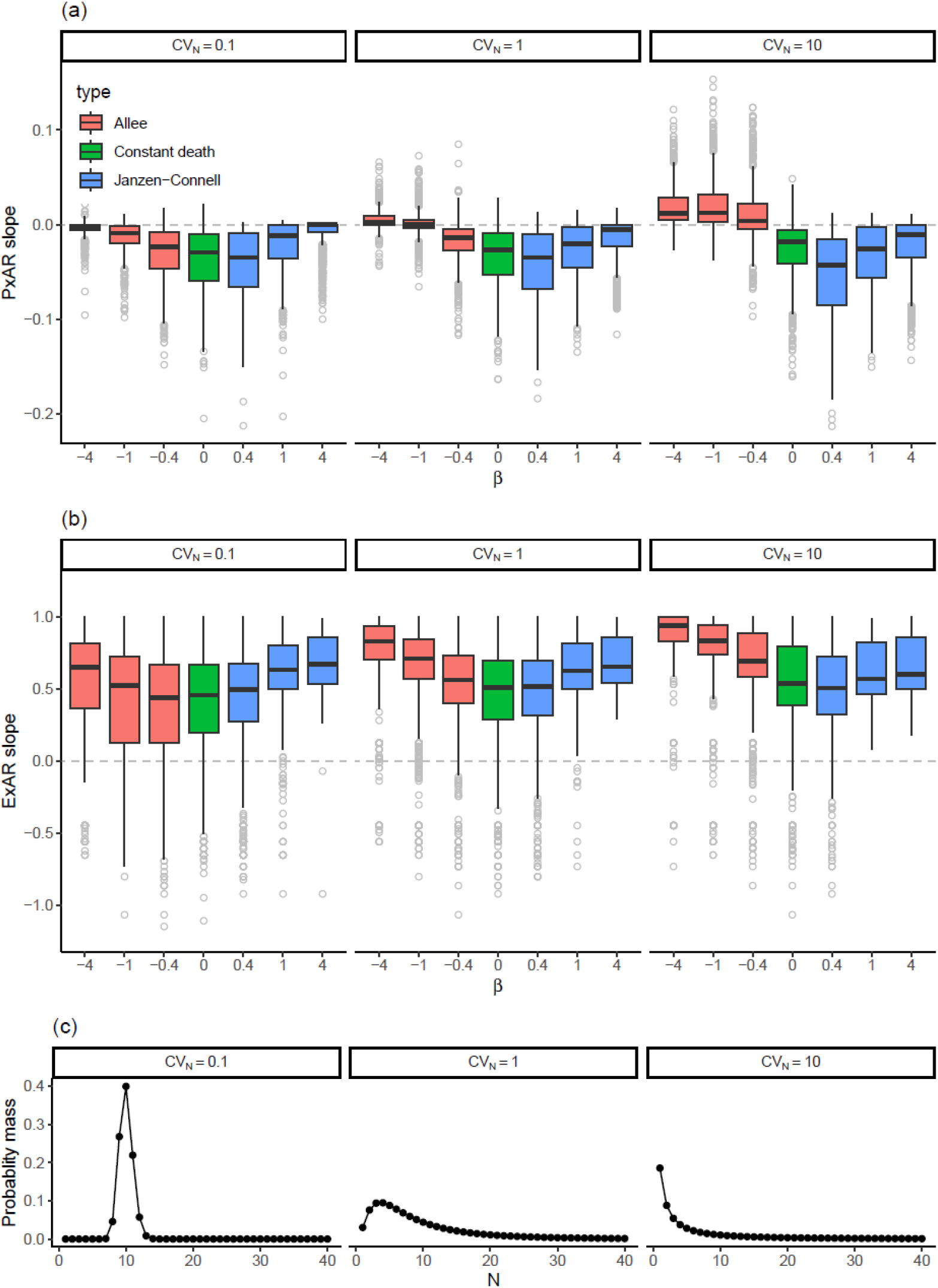
The effect of the sign and magnitude of density-dependence of death rate (determined by parameter *β*, Table 2) on the slope of PxAR (a) and ExAR (b) in point pattern simulations. Panels are divided according to three levels of the *CV*_*N*_ parameter, which affects the shapes of the regional species-abundance distribution (SAD), from more more even (left) to uneven (right). These SADs are illustrated in panel (c).

**Figure 5.**
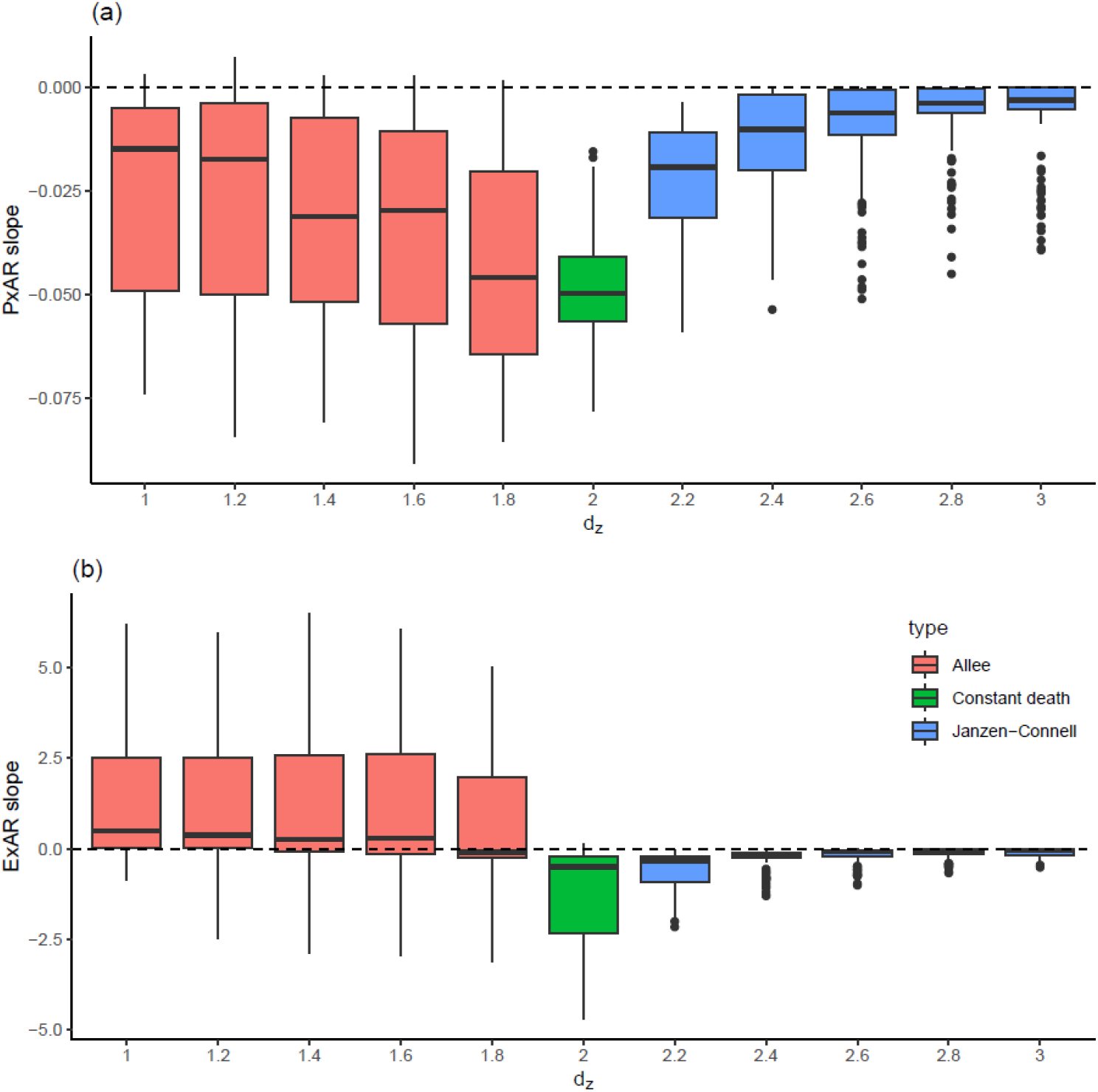
The effect of parameter *d*_*z*_ on the slope of PxAR (a) and ExAR (b) in Lotka-Volterra simulations. The parameter *d*_*z*_ regulates the sign and magnitude of the density dependence of death rate.

### Hypothesis 2

We found that Janzen-Connell effect in point pattern simulations produced negative slopes of PxAR (Fig. 4a, blue boxes), consistent with Hypothesis 2. This was more pronounced when evenness of the regional SAD (parameter *CV*_*N*_) was low; in such cases, the slope of the PxAR was more negative than a slope expected from constant *P*_*death*_ (green box in Fig. 4a when *CV*_*N*_ *= 10*). In all other cases in the point pattern, the PxAR slope under the Janzen-Connell effect was similar to the slope of the constant *P*_*death*_. In the Lotka-Volterra simulations, however, the PxAR slope was negative (as expected under Hypothesis 2), but higher than in the constant *P*_*death*_ scenario (Fig. 5a). In both simulation models, the stronger was the negative density dependence (the more parameters *β*and *d*_*z*_ increased), the more the PxAR slope approached 0 (Fig. 5a).

### Hypothesis 3

As expected, ExAR was affected by different parameters than was the PxAR (Fig. 3). In line with Hypothesis 3, the total number of species (*S*_*frac*_ and *S*) had a stronger effect on ExAR than on PxAR in both point pattern and Lotka-Volterra simulations. In addition, in the Point pattern simulations (Fig. 3a, b), the ExAR (but not PxAR) slope was strongly affected by the evenness of the regional SAD (parameter *CV*_*N*_) and by con-specific spatial aggregation (parameter *σ*), also in line with Hypothesis 3.

## 5 Discussion

To get back to our title question: Should regional species loss be faster, or slower, than local loss? Our main finding is that this depends on the strength of the relationship between population density (rarity) and per-individual probability of death. This role of rarity is even more important than other drivers of the extinction scaling, such as the conspecific aggregation. Particularly the latter was pointed out as an important driver of extinction scaling by (Yan et al. 2022) We confirm Yan et al.’s (2022) results, but we also extend them by adding an even more important mechanistic driver than a simple conspecific aggregation.

Another important finding is the difference between the spatially explicit point pattern simulations and the Lotka-Volterra simulations. There was a general scarcity of the positive PxAR (lower local and higher regional per-species extinction rates), which we only invoked in the point pattern simulations, particularly in uneven initial species-abundance distributions (SAD). In other words, although Hypothesis 2 was generally confirmed, Hypothesis 1 was confirmed only in the point pattern simulations. Here is our explanation: With the point patterns, we simulated a one-time loss event, in which rare or common species were present at time 1, and the Allee or Janzen-Connell effects could target these rare or common species, leading to extinction scaling that was in line with our hypotheses. However, the Lotka-Volterra simulations represented a system at a steady state where rare or common species were constantly targeted in every step of the simulation, thus leading to an elimination of rarity or commonness from the system, making the SAD more even, and the PxAR negative.

We thus expect Hypotheses 1 and 2 to apply particularly after irregular, exceptional, or one-time mortality events. Examples are abrupt human-caused changes in landscape management, land cover, disturbances such as fires, or extreme climate events. However, in stable systems closer to a steady state where mortality is a continuous process, we expect a generally negative PxAR, irrespectively to whether Allee or Janzen-Connell effects take place. This is also what has been observed in such systems, for example in a long-term time series of Czech birds, or in the Barro-Colorado forest plot (Keil et al. 2018).

We can turn this insight to broad predictions. For example, in birds, we know that most global extinctions concern endemic species from islands (Loehle and Eschenbach 2012), and most of these are due to anthropogenic pressures, particularly due to the spread of invasive species such as domestic cats; in the grand scheme of things, this is a one-time disturbance. We would thus expect this to cause a positive global PxAR, with high global and low local per-species probability of extinction.

Concerning the total number of extinct species, and in line with Hypothesis 3, the direction of the ExAR was affected by other important drivers, i.e. other than by the relationship between per-individual death rate and rarity, specifically by factors affecting the species-area relationship. As a consequence, the observed slopes of ExAR were variable, and not correlated with PxAR slopes. This is also what has been broadly observed in empirical data where both negative and positive ExARs are common (Keil et al. 2018).

Some may argue that our exercise only involved limited numbers of individuals, which may correspond to a small spatial extent. Over regional and continental extents, total abundances of real-world species are likely orders of magnitude higher. However, we tentatively suggest that our results are still relevant for large extent situations involving species’ geographic ranges; one simply can replace point occurrence for occupied grid cells across a continent, and one can replace per-individual probability of death by per-grid cell probability of loss of the species per grid cell. Similar principles of extinction scaling should then apply both for grid cell occupancy and our point pattern simulations. Moreover, once we consider occupancy, the same principles also apply for organisms for which individuals are not well defined, such as some plants. This connection between presence and absence of individuals, abundance, and species occupancy across scales, has been described in the form of occupancy-area relationship (Kunin 1998).

In our simulations, we assumed that the Allee or Janzen-Connell effects apply to all species in the community, while real-world species likely differ in susceptibility to these effects, or both types of effect can operate within a single system. We suspect that the latter may even lead to non-linear (hump-shaped, U-shaped) PxAR and ExAR, as observed in real-world data, and something that we also miss in our analyses as we only fit linear functions. This can be investigated in the future.

We envision several follow-ups: First, a similar exercise can be done for the rates of species gain. Recently, Leroy et al. (Ecography, *in press*) proposed a simple idea that distinguishes local gain caused by within-regional homogenization from gain caused by colonization from an outside pool, and how this affects the spatial scaling of species gain. A quantitative evaluation of such a proposition, in the spirit of the simulations presented here, would be a logical next step. Second, a mechanism could be tested that also includes spatial inter-specific dependence of probability of death among individuals. This can either be caused by spatially auto-correlated mortality events such as fires, floods, or pest outbreaks, or by loss of species that maintain vital mutually beneficial interactions. A loss of one species can thus lead to a domino effect in which other species are lost, maybe with interesting consequences for the scaling of the loss.

In all, we see the main value of this exercise in the strengthening of the argument that biodiversity loss, and biodiversity change in general, must be considered as a strongly scale-dependent problem. This has so far been acknowledged based on numerous empirical observations (Keil et al. 2011, 2018, Powell et al. 2013, Chase et al. 2019, Leroy et al. 2023). Here, by linking individual demographic parameters to the community level, we add a direct ecological mechanistic interpretation of this scale dependence.

## 6 Acknowledgements

PK and FL were funded by the European Union (ERC, BEAST, 101044740). Views and opinions expressed are however those of the author(s) only and do not necessarily reflect those of the European Union or the European Research Council Executive Agency. Neither the European Union nor the granting authority can be held responsible for them. We are thankful to William E. Kunin for a discussion that raised the issue of independence among species.

## 7 Author contributions

PK and FL conceived the ideas. ATC designed the Lotka-Volterra simulations. VB designed the Barták function. PK performed and analyzed the simulations and led the writing, with contributions from all co-authors.

## Notes

https://github.com/petrkeil/extinction_scaling_simulations

### Competing Interest Statement

The authors have declared no competing interest.

